# Evolution of brilliant iridescent feather nanostructures

**DOI:** 10.1101/2021.05.31.446390

**Authors:** Klara K. Nordén, Chad M. Eliason, Mary Caswell Stoddard

## Abstract

The brilliant iridescent plumage of birds creates some of the most stunning color displays known in the natural world. Iridescent plumage colors are produced by nanostructures in feathers and have evolved in a wide variety of birds. The building blocks of these structures—melanosomes (melanin-filled organelles)—come in a variety of forms, yet how these different forms contribute to color production across birds remains unclear. Here, we leverage evolutionary analyses, optical simulations and reflectance spectrophotometry to uncover general principles that govern the production of brilliant iridescence. We find that a key feature that unites all melanosome forms in brilliant iridescent structures is thin melanin layers. Birds have achieved this in multiple ways: by decreasing the size of the melanosome directly, by hollowing out the interior, or by flattening the melanosome into a platelet. The evolution of thin melanin layers unlocks color-producing possibilities, more than doubling the range of colors that can be produced with a thick melanin layer and simultaneously increasing brightness. We discuss the implications of these findings for the evolution of iridescent structures in birds and propose two evolutionary paths to brilliant iridescence.

## Introduction

Many animal colors—and indeed some plant, algae and possibly fungus colors (Brodie et al., 2021)—are structural, produced by the interaction of light with micro- and nano-scale structures (reviewed in Kinoshita et al., 2008). In birds, structural colors greatly expand—relative to pigment-based mechanisms—the range of colors birds can produce with their feathers (Stoddard and Prum, 2008). Some structural colors are iridescent: the perceived hue changes with viewing or lighting angle. Iridescent coloration features prominently in the dynamic courtship displays of many bird species, including birds-of-paradise (Paradisaeidae), hummingbirds (Trochilidae) and pheasants (Phasianidae) (Greenewalt et al., 1960; Stavenga et al., 2015; Zi et al., 2003). These dazzling displays showcase the kind of bright and saturated iridescent colors that have previously been qualitatively categorized as “luxurious” (Auber, 1957) or “brilliant” (Durrer, 1977), in contrast to the more muted “faint” (Auber, 1957) or “weak” (Durrer, 1977) iridescent colors of, for example, a brown-headed cowbird. Following these authors, we use the terms “weak” and “brilliant” to describe this difference in color appearance, where brilliant iridescence describes colors of high saturation and brightness, and weak iridescence colors of low saturation and brightness. Typically, brilliant iridescence is associated with more complex feather nanostructures than weak iridescence. All iridescent feather coloration is produced by nanostructures in the feather barbules consisting of melanin-filled organelles (melanosomes) and keratin (Figure 1), but brilliant iridescent coloration arises from light interference by photonic crystal-like structures (henceforth photonic crystals), while weak iridescent coloration is produced by structures with a single layer of melanosomes (Durrer, 1977). A photonic crystal is defined by having periodic changes in refractive index (Joannopoulos et al., 2008); in feather barbules, this is created by periodic arrangements of melanosomes in keratin. By adding more reflection interfaces, a photonic crystal greatly amplifies color saturation and brightness compared to a single-layered structure, the latter which functions as a simple thin-film (Kinoshita et al., 2008).

**Figure 1.**
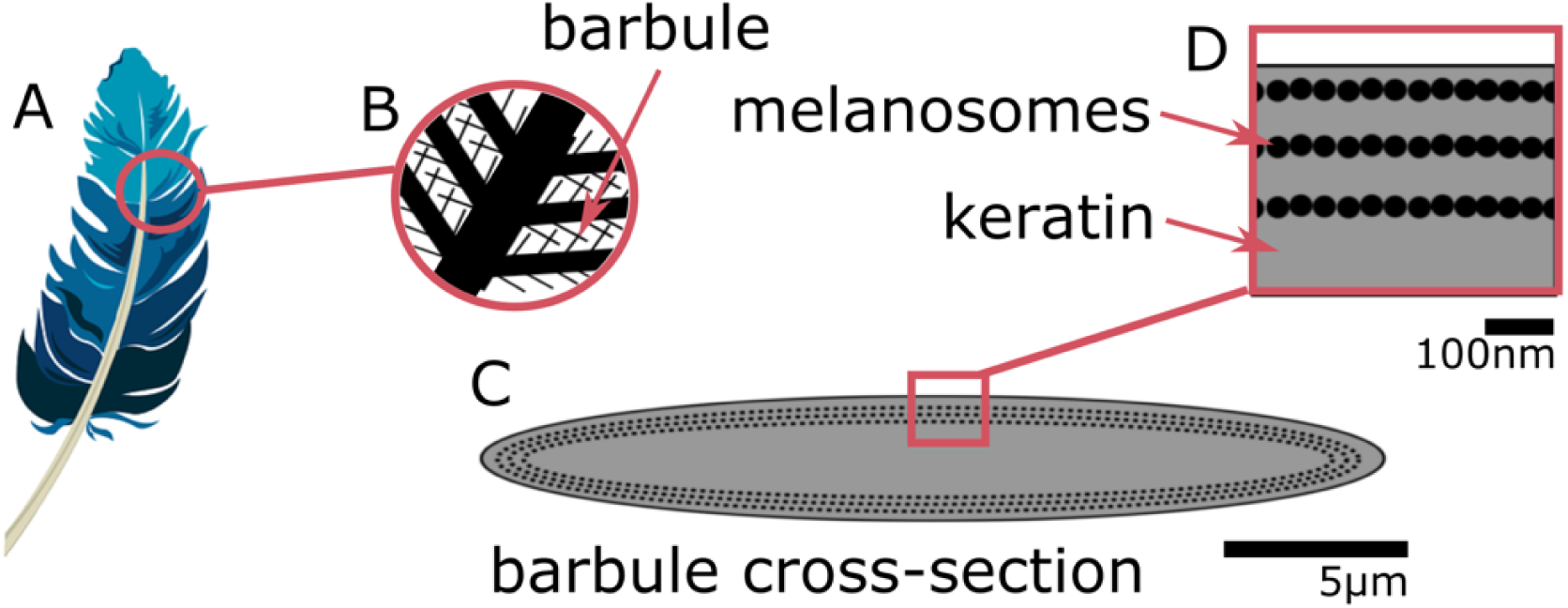
Iridescent plumage is produced by nanostructures in the feather barbules. A vaned feather (A) consists of branching structures where the barbules (B) are the interlocking filaments. A cross-section of a barbule from an iridescent feather (C) reveals the intricate nanostructure responsible for the color, consisting of layers of melanosomes in keratin (D). Blue feather in A from Pixabay, licensed under Pixabay License (full details in Supp. Note 1).

In iridescent feathers, it is not just the arrangement of melanosomes that can vary: the melanosomes also come in a variety of different shapes. Durrer, 1977 classified melanosomes into five main types: 1) thick solid rods (S-type, Figure 2A); 2) thin solid rods (St-type, Figure 2B); 3) hollow rods (with an air-filled interior, R-type, Figure 2C); 4) platelets (P-type, Figure 2D); and 5) hollow platelets (K-type, Figure 2E). Thick solid rods are typically found in single-layered structures producing weak iridescence (Figure 2A), while thin solid rods, hollow rods, platelets and hollow platelets occur in photonic crystals producing brilliant iridescence (Fig 2B-D). This diversity is extraordinary given that the shape of melanosomes in other melanized vertebrate tissues, including black and grey feathers, is typically a solid rod (D’Alba and Shawkey, 2019). The thick solid rods found in weakly iridescent feathers resemble the melanosomes found in plain black feathers (Durrer, 1977) and are likely ancestral to the four more elaborate, derived melanosome shapes (Shawkey et al. 2006; Maia et al. 2012). Because the derived melanosome shapes (but not the ancestral thick solid rods) are arranged as photonic crystals, these two innovations together—novel shapes and photonic crystal structure—may have been critical for the evolution of brilliant iridescence. Supporting this idea, Maia et al., 2013b showed that the evolution of hollow and/or platelet-shaped melanosomes in African starlings (Sturnidae) was associated with great expansions in color diversity and increases in brilliance. Moreover, Eliason et al., 2013 used optical modeling and plumage color measurements of the violet-backed starling (*Cinnyricinclus leucogaster*) and wild turkey (*Meleagris gallopavo*) to show that hollow rods increase the brightness of iridescent colors compared to structures with solid rods.

**Figure 2.**
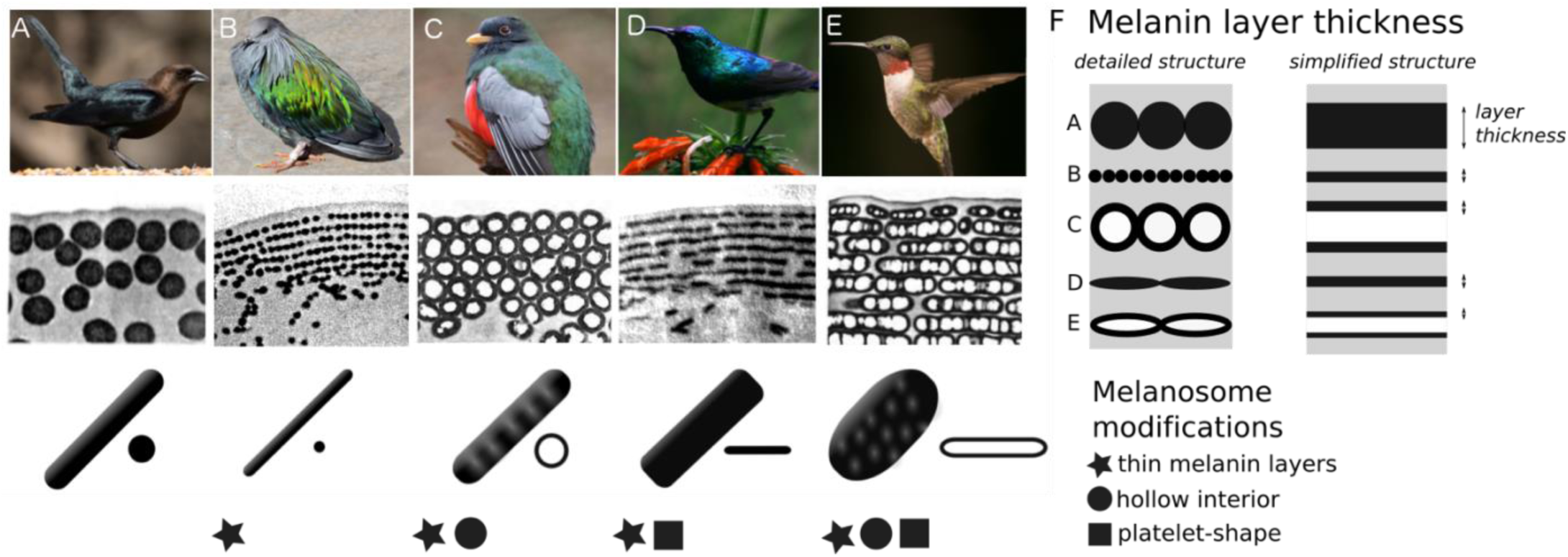
Iridescent feather nanostructures are diverse. Structures can vary both in melanosome type and melanosome organization. There are five main types of melanosomes (shown as schematics in bottom row, A-E) and two main types of structural organization (shown by microscope images of barbule cross-sections, middle row: single-layered (A) and photonic crystal (B-E)). A single-layered structure with thick solid rods (A) gives rise to the dark, black-blue iridescence of a brown-headed cowbird (*Molothrus ater*). This type of structure generally gives rise to “weak” iridescent colors, with low color saturation and brightness. Photonic crystals (B-E) with multiple layers of melanosomes generally give rise to “brilliant” iridescent colors, with high saturation and brightness. Thin solid rods (B) in a multilayer configuration (also called a one-dimensional photonic crystal) produce the iridescent colors of the Nicobar pigeon (*Caloenas nicobarica*). In the elegant trogon (*Trogon elegans*), the iridescent green color is produced by hexagonally packed hollow rods (C). Sunbird (here the variable sunbird, *Cinnyris venustus*) barbules contain melanosomes stacked in multilayers, with solid platelet-shaped melanosomes serving as the building blocks (D). The fifth melanosome type is a hollow platelet (E), which forms multilayer configurations in many hummingbird species (here a ruby-throated hummingbird*, Archilochus colubris*). The five types of melanosomes are characterized by different combinations of three key modifications: thin melanin layers, hollowness and platelet shape, which are indicated as symbols under each melanosome type. Thin melanin layers are present in four melanosome types, but they are achieved in different ways, as is shown by the schematic in F. A simplified diagram of each melanosome type (F, right) shows how solid forms translate to a single melanin layer, while hollow forms create two thinner melanin layers intersected by an air-layer. All photographs (top row) are under a Public Domain License (details in Supp. Note 2). Transmission Electron Microscope images from Durrer (1977), reproduced with permission.

While previous studies focusing on nanostructural evolution and color-producing mechanisms in a variety of avian subclades (Eliason et al., 2020, 2015, 2013; Eliason and Shawkey, 2012; Gammie, 2013; Gruson et al., 2019; Maia et al., 2013b; Quintero and Espinosa de los Monteros, 2011) have given us valuable insights into the evolution and optics of iridescent structures, they have focused on specific species, subclades or a particular melanosome type in isolation. Thus, they have not uncovered the broader, general principles governing the evolution of brilliant iridescent plumage, and several key questions remain unanswered.

Why have bird species with brilliant iridescence evolved not one but four different melanosome types? How are these melanosome types phylogenetically distributed? Are particular melanosome types associated with different plumage colors? Since Durrer’s publication in 1977, there has been no broad-scale evolutionary analysis of the melanosomes in iridescent feathers, and no study has compared the optical effects of all five of Durrer’s melanosome types. To find general principles underlying differences in color production, we identify key modifications of each melanosome type that based on optical theory are likely to be important. This enables us to compare the five melanosome types rigorously, since each type can have several modifications. For example, a hollow platelet (Figure 2E) has both an air-filled interior and a flattened shape, both of which might influence feather color—perhaps in different ways. Therefore, a simple comparison of the melanosome types cannot reveal which modifications are contributing to differences in color production and how.

In this study, we search for general design principles underlying the production of brilliant iridescent coloration. First, we identify three key modifications of melanosomes in brilliant iridescent structures: thin melanin layers, hollowness, and platelet shape, (Figure 2). Second, we create a feather iridescence database using published descriptions of iridescent feather structures. Using the database, we explore the evolutionary history of the three key modifications of brilliant iridescent structures. Third, we use optical modeling to simulate colors that could be produced with each melanosome type; we estimate light reflectance from 4500 different structures using parameter ranges derived from the database. Finally, we analyze spectral data from 120 plumage regions across 80 diverse bird species with known nanostructures to test the predictions of our optical model.

## Results

### 1) Identifying key melanosome modifications

The size, composition and shape of materials that form the periodic layers in a photonic crystal can all contribute to its reflectance properties (Joannopoulos et al., 2008). In iridescent structural feather colors, the layers are formed by melanosomes, and we can identify three melanosome modifications that likely have important optical effects. We define these modifications relative to the thick solid rods found in weakly iridescent feathers, since we presume these to be “unmodified” or “minimally modified” from melanosomes found in other non-iridescent melanized tissues, which they closely resemble. The three modifications are: thin melanin layers (size of layers), an air-filled interior (layer material composition), and platelet shape (shape of layers). “Thin” here refers to something thinner than the ancestral thick solid rods. A “melanin layer” refers to a single layer in the optical structure. For solid rods and platelets, this is simply the rod or platelet diameter, but for hollow rods and platelets, this is thickness of a single melanin wall (Figure 2F). Each of Durrer’s five melanosome types can be described in terms of the absence/presence of one or several modifications (Figure 2).

What are the potential optical advantages of melanosomes with these features? First let us consider thin melanin layers. Thin melanin layers may tune the structure so that it reflects optimally in the bird-visible spectrum. This possibility was raised by Durrer, 1977, who noted a convergence towards thin [melanin] layers in structures producing brilliant iridescent colors. However, his work is only available in German, and this idea has remained largely overlooked. We refine and extend his idea here using established optical theory, specifically multilayer optics (reviewed in Kinoshita 2008; Kinoshita et al. 2008). To produce first order interference peaks, which will result in brighter colors than higher order interference peaks, the optical thickness (thickness×refractive index) of each repeating unit in a one-dimensional photonic crystal (also often termed multilayer, Figure 2B, D-E) should approximate half a wavelength 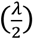 (cf. Durrer, 1977; Kinoshita et al., 2008; Land, 1972). The repeating unit in an iridescent feather nanostructure consists of one layer of melanosomes and one layer of keratin, and we can therefore express this as 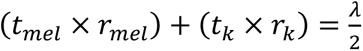, where *t* is the thickness and *r* is the refractive index of melanin (*mel*) and keratin (*k*) layers. Among the configurations that satisfy this condition, maximum reflection is achieved when both layers have equal optical thickness, which can be expressed as 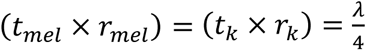 (Kinoshita et al., 2008; Land, 1972). From this, we can calculate the range within which we would expect melanin optical layer thickness to fall: 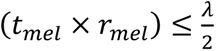, with maximum reflectance at 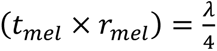. In practice, this means we should expect to see melanin layer thicknesses of at most 75nm (respectively 206nm) with maximum reflectance at 38nm (respectively 103nm) for the ends of the bird-visible spectrum; 300nm (respectively 700nm). Here, we use the refractive indices *r_mel_* = 2 for 300nm and *r_mel_* = 1.7 for 700nm, obtained by Stavenga et al., 2015. The typical diameter of melanosomes found vertebrates is much larger than this range, ∼300nm (Li et al., 2014).

Now let us consider why melanosomes with hollow interiors and/or platelet shapes might be advantageous. A hollow interior could increase reflection by creating a sharper contrast in refractive index in the structure (Durrer, 1977; Eliason et al., 2013; Kinoshita, 2008; Land, 1972; Stavenga et al., 2018), while platelet-shaped melanosomes could increase reflection by creating smooth, mirror-like reflection surfaces (Durrer, 1977; Land, 1972). Moreover, the thin platelet shape might allow for more layers to be packed within a photonic crystal, which would increase total reflection (Maia et al., 2013b).

Which of the four derived melanosome types in brilliant iridescent feathers possess these modifications? Hollowness and platelet shape are each present in two types, but thin melanin layers are likely shared by all four derived melanosome types. This was indicated by Durrer, 1977, but he never analyzed the distribution of layer thickness among the structures he measured. However, this potential convergence hints at the intriguing possibility that all four derived types present diverse paths to the same end: achieving optimal melanin layer thickness. A hollow interior or a platelet shape may simply be different mechanisms for reducing melanin layer thickness. This would also explain why thick solid rods are typically only found in single-layered structures. Single-layered structures typically function as thin-films, where only the overlying keratin cortex produces the interference colors (Doucet, 2006; Lee et al., 2012; Maia et al., 2009; Yin et al., 2006). The layer of melanosomes only functions to delimit the keratin layer, therefore the thickness of the melanin layer itself is irrelevant. Thus, there would be no selection pressure to decrease melanin layer thickness in single-layered structures, and we would expect the ancestral condition (thick solid rods) to remain.

We suggest that the diverse melanosome types found in brilliant iridescent structures evolved to generate thin melanin layers in different ways. This possibility has not been investigated previously, probably because melanosome types are generally analyzed on the basis of their overall morphology rather than—as we have proposed here—on the basis of specific optical modifications.

### 2) Evolution of modified melanosomes in iridescent structures

We surveyed the literature for all published descriptions of iridescent feather structures in order to build a species-level database (henceforth the feather iridescence database) of key structural parameters (Figure 8). These parameters included melanosome type (solid rod, hollow rod, solid platelet, hollow platelet), melanin layer thickness, details about the structure (single-layered or photonic crystal), and size of the internal air pockets. We found that iridescent feather nanostructures have been described in 306 bird species representing 15 different orders and 35 families. The feather iridescence database, which includes a complete list of the references we consulted, is included in the Supplementary Information.

Descriptions of iridescent feather structures are taxonomically biased, with some groups well represented (>20 species represented in the database: Sturnidae, Trochilidae, Phasianidae, Trogonidae and Anatidae) but most groups sparsely sampled (<5 species in the database) or absent (e.g., Picidae). Even in well-sampled groups (e.g., Trochilidae), the feather structures of only about 15% of all the species in the family have been described. Some published descriptions included measurements of every structural parameter, while others only included partial information on melanosome modifications. For 61% of the species has the thickness of melanin layers been described, while almost all species have complete information on the presence/absence of melanosome hollowness and/or platelet shape (92%). Most species records (83%) described the type of structure (single-layered or photonic crystal). These data, though taxonomically biased, allowed us to describe the properties of the three melanosome modifications we defined (thin melanin layers, hollowness, platelet shape). Using an avian phylogeny (Jetz et al., 2012), we mapped these modifications for all 280 species for which complete information on melanosome type was present in our database (Figure 3A). Although these species represent only a fraction of those with iridescent feathers, the major iridescent orders are represented. Our analysis thus provides a broad snapshot of iridescent feather structure diversity and evolution across birds. In the sections below, we use this dataset to test functional hypotheses for each modification and discuss their evolutionary pattern in more detail.

**Figure 3.**
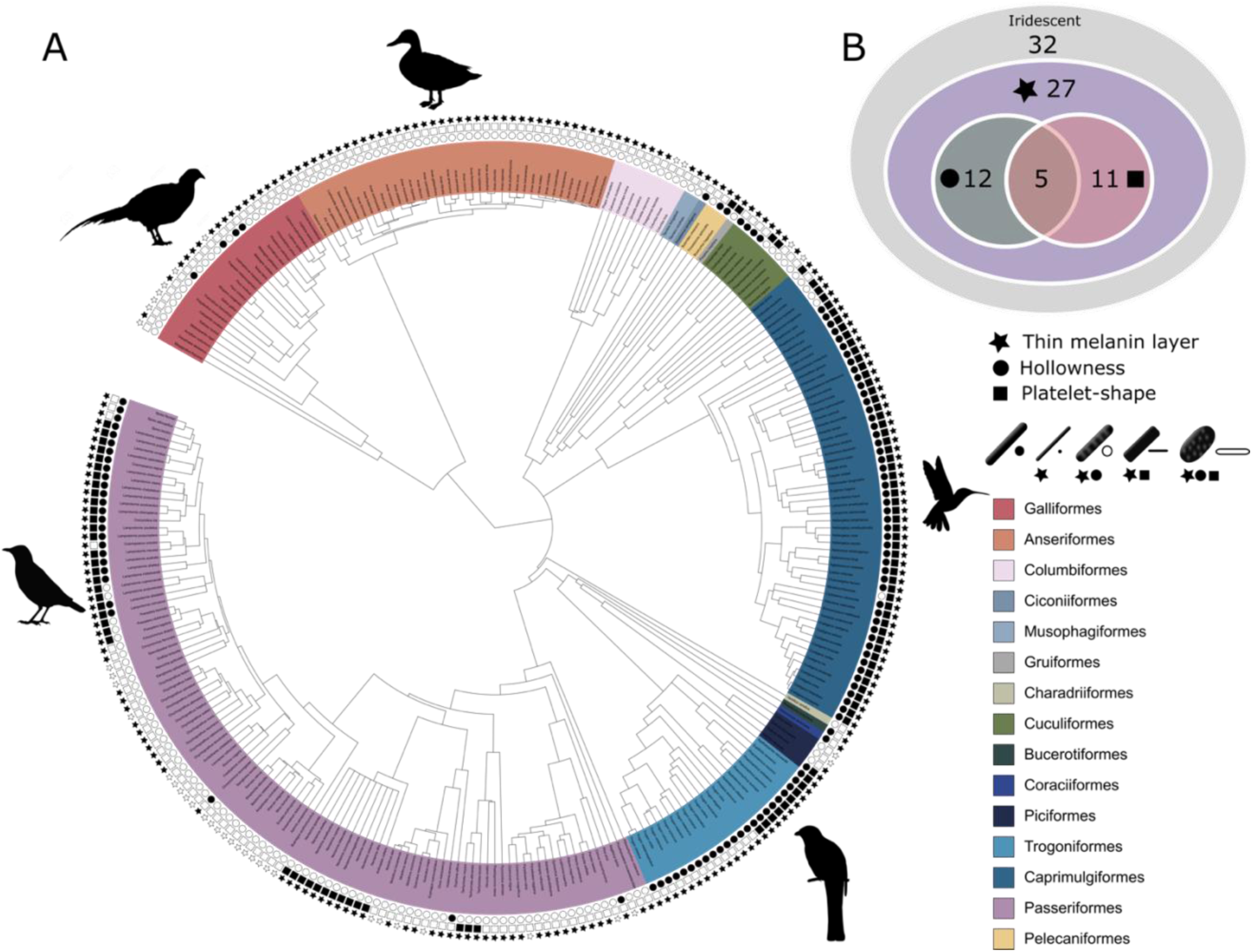
Evolutionary distribution of three key melanosome modifications in iridescent structures: thin melanin layers (star), hollowness (circle) and platelet shape (square). Schematics of melanosomes in the key show how combinations of modifications correspond to each melanosome type. A) Melanosome modifications mapped to a phylogeny including all species in the feather iridescence database (280 species, 26 species lacking data on melanosome type excluded). Note that where data on melanin layer thickness was not available for a species with hollow and/or platelet-shaped melanosomes, they were assumed to have thin melanin layers, since all known structures do. Silhouettes shown for the five families that are most well represented in the feather iridescence database (>20 species represented in the database): Sturnidae, Phasanidae, Anatidae, Trogonidae and Trochilidae. B) Venn diagram showing the number of bird families in the feather iridescence database where each modification was present. The majority of bird families with iridescent plumage studied have evolved thin melanin layers, and there are no hollow or platelet-shaped melanosomes which have not also evolved this modification. A similar number of families have hollow or platelet-shaped melanosomes, but only five families have evolved both modifications together. Note that this plot depicts the number of occurrences of each modification, not independent evolutionary origins. Silhouettes from Phylopic.org, licensed under a Public Domain License (full details in Supp. Note 3).

#### Thin melanin layers

We have suggested that all four melanosome types found in brilliant iridescent structures (Figure 2B-E) share a common trait: a reduction in melanin layer thickness. This is plausible based on the measurements and description of melanosome types given by Durrer, 1977 but has not been formally quantified. In fact, Durrer’s division of solid rods into a thinner (diameter of ∼100nm) and thicker variety (diameter of ∼200nm) has not been previously tested or precisely defined. In the current literature, solid rods are often treated as a single type with a continuum of diameters (Eliason et al., 2013; Maia et al., 2013b; Nordén et al., 2019). Thus, to study the evolution of thin melanin layers, we first needed to define this trait using the feather iridescence database. Specifically, we used the feather iridescence database to show that: 1) Solid rods can be divided into two distinct distributions (a thinner and thicker variety), 2) Hollow and/or platelet shaped melanosomes have equally or thinner melanin layers than thin solid rods, demonstrating that they share this modification.

Exploring the distribution of melanosome diameter in all solid rods, we found a significant bimodal distribution (Figure 4, unimodality rejected, p<0.001, bimodality not rejected, p=0.86). Based on the bimodal distribution of melanosome diameters in solid rods, we define “thick solid rods” as those with a diameter >190nm and “thin solid rods” as those with a diameter ≤190nm. It should be noted that this definition differs slightly from Durrer’s categorization, who notes a range of 70-140nm for the thin solid rods he measured (Durrer, 1977). It can be seen that thick solid rods overlap considerably with melanosomes in black feather (data from Li et al., 2012), supporting the hypothesis that thick solid rods represent minimally or unmodified melanosomes. Iridescent structures most likely evolved from black plumage (Maia et al., 2012; Shawkey et al., 2006), therefore we can use the size of melanosomes in black melanosomes to represent an “unmodified” melanosome. In contrast, there is no overlap between the melanin thickness of melanosomes in black feathers and thin solid rods, suggesting that this is a considerable modification from the ancestral state.

**Figure 4.**
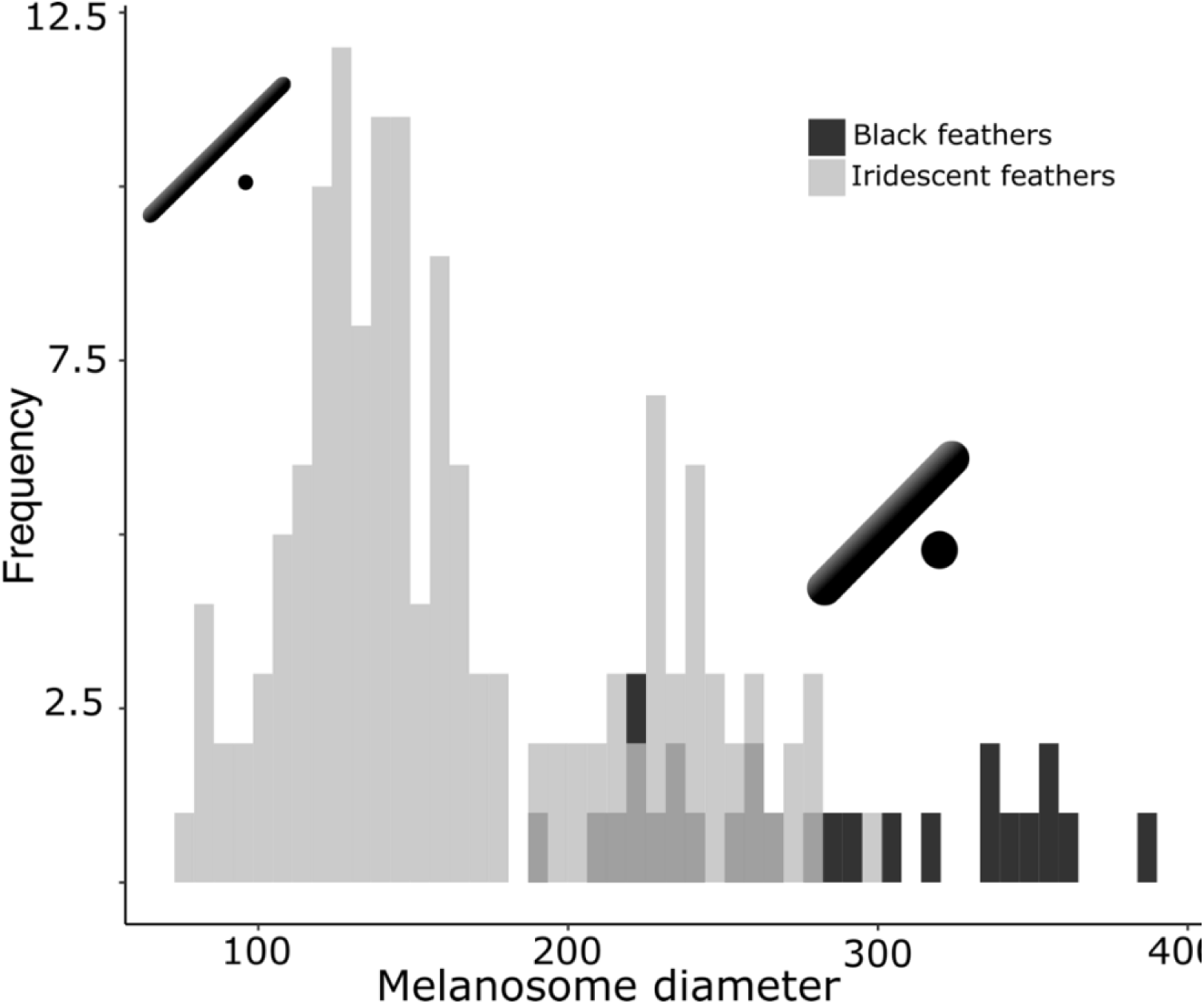
There are two distinct types of solid rods in iridescent structures: thick solid rods and thin solid rods. This is evident from the clear bimodal distribution shown by the histogram of melanosome diameters found among solid rods in the feather iridescence database (grey). Based on this distribution, we define “thin rods” as any solid rod with a diameter ≤190nm. Plotted in black is the distribution of diameters from melanosomes in black feathers (data from Li et al. 2012), which overlaps with the distribution of thick solid rods in iridescent structures.

We can now define “thin melanin layers” as any melanosome with melanin layers ≤190nm. Using this new definition, we found that all hollow and/or platelet-shaped melanosomes can indeed be classified as having thin melanin layers (range 24-139nm, Figure 5). Whether a single melanin wall in hollow melanosomes always represent one melanin layer is debatable: some photonic crystals with hollow melanosomes have little or no keratin interspersed between melanosome layers (e.g., Figure 2C, E). In these cases, it may be more appropriate to think of a single melanin layer as the sum of two melanin walls. However, all hollow forms in photonic crystals have a melanin wall thickness of <100nm (Figure 5), so they would still qualify as “thin”. All four derived melanosomes with thin melanin layers have significantly thinner melanin layers than melanosomes in black feathers and thick solid rods (phylogenetic pairwise t-test, all p<0.01, see details in Table S1).

**Figure 5.**
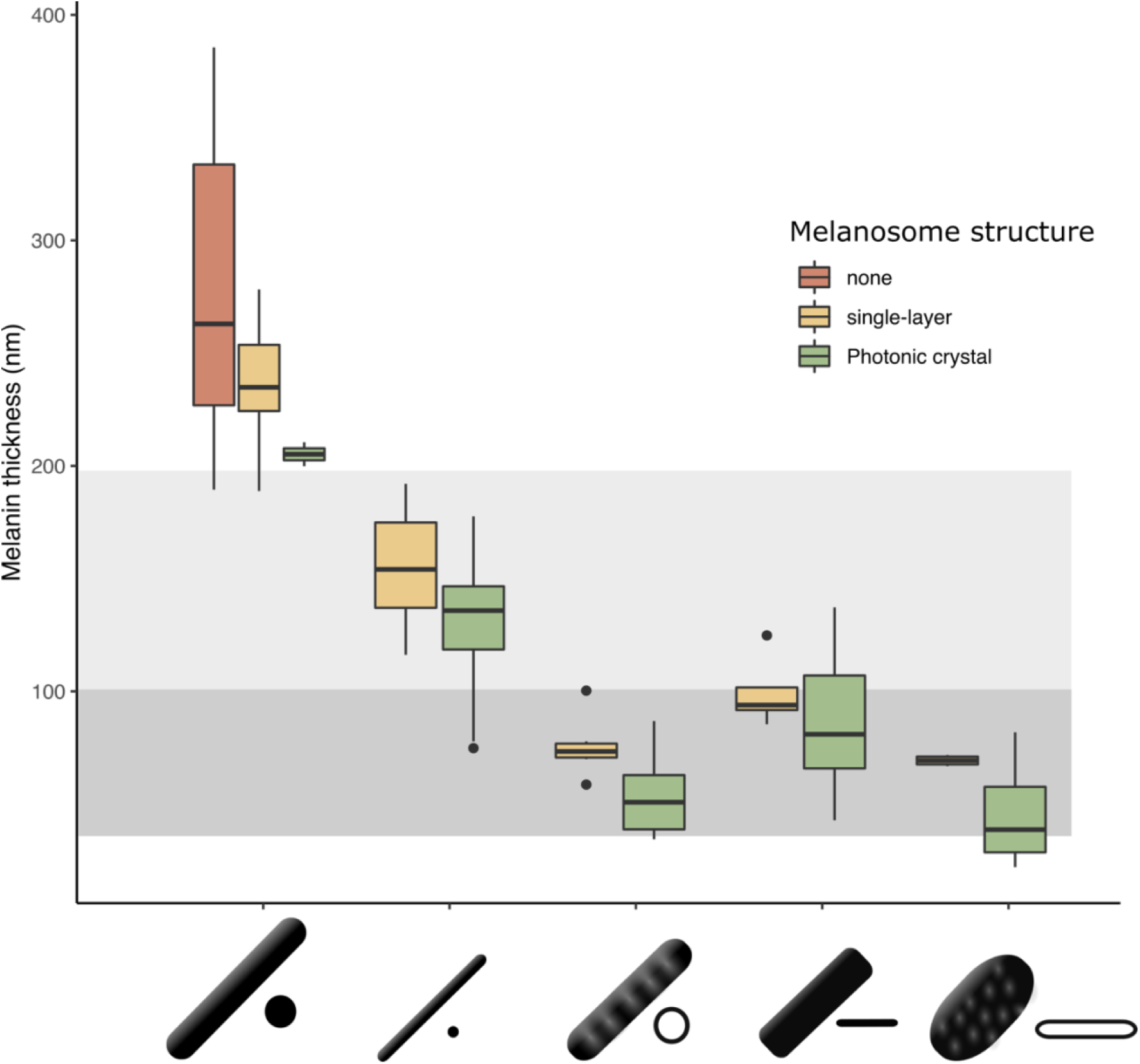
The thickness of melanin layers in brilliant iridescent structures has converged towards the theoretical optimal range, where optical thickness equals 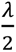 (light grey box, for bird-visible spectrum). Boxplot shows the distribution of melanin layer thickness for each melanosome type in single-layered structures (yellow) and photonic crystals (green) in the feather iridescence database. “None” corresponds to melanosomes in a black feather without organization. All melanosome types except thick solid rods, which are predominantly found in single-layered structures with weak iridescence, have converged towards an optical thickness of 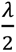. Hollow and platelet forms often reach thicknesses closer to 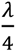, which can in theory form ideal multilayers (dark grey box, for bird-visible spectrum).

Next, we tested our hypothesis that thin melanin layers evolved for a specific optical benefit—to allow photonic crystals to produce bright and saturated colors. We have already shown that the four derived melanosomes share a modification for thin melanin layers, but it is possible that this evolved for reasons unrelated to color production, such as to minimize the cost of melanin production. We predicted that if thin melanin layers did evolve for the optical benefit, they should have converged on the optimal range for producing bright interference peaks in the bird-visible spectrum (38-206nm). In addition, we predicted that melanosomes with a thickness outside this favorable range should be rare or absent in photonic crystals. We found that all melanosomes with a decreased melanin layer thickness indeed have converged on thicknesses well within this optimal range (Figure 5). Moreover, all derived melanosome types achieve thicknesses of 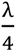 (38-103nm). Such structures could in theory produce ideal multilayers, which produces the greatest reflectance for a two-material reflector (Land, 1972). We also found that the great majority of photonic crystals contain melanosomes with thin melanin layers (98%). Overall, these findings are compatible with the hypothesis that the primary benefit of thin melanin layers is to produce bright and saturated colors.

The importance of this modification for iridescent color production can also be inferred from its phylogenetic distribution. Over 80% of all families represented in the feather iridescence database have evolved thin melanin layers (27 out of 32 families, Figure 3B). The families that lack the thin modification also lack species with brilliant iridescent plumage (Numididae, Aegithinidae, Irenidae, Buphagidae, Megapodiidae and Lybiidae).

#### Hollowness

Hollowness occurs in both rod-shaped and platelet-shaped melanosomes. However, whether the size of internal air pockets differs in rods compared to platelets has never been tested. If air pockets function primarily in producing strong interference colors in bird-visible wavelengths, we predict that there should be no difference between air pocket diameter in rods and platelets and that diameters should be constrained between 75-175nm (n_air_ = 1) to give similar optical thicknesses as melanin layers 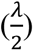, thereby optimizing reflection (see Results, §1). On the other hand, if hollowness evolved for different reasons in rods and platelets, and/or for non-optical functions, they may differ. Air pocket size (length in the shortest dimension, see Figure 8) ranged from 50-251 nm and did not differ significantly between rods and platelets (phylogenetic ANOVA, F(1, 55)= 16.8, p=0.176, df=3). This range does indeed include the thickness that would produce interference colors of the first order in the bird-visible range. Together with our results on melanin layer thickness, this indicates both air pockets in hollow melanosomes and thin melanin layers are simultaneously tuned to produce bright and saturated colors.

Our phylogenetic analysis shows that a hollow interior has evolved in at least 12 bird families, or 34% of all families in the feather iridescence database (Figure 3B). Many families with brilliant iridescence are included, such as Phasianidae, Trochilidae and Sturnidae. However, hollow melanosomes do not appear to be a requirement for brilliant iridescence. Unlike thin melanin layers, which are present in all families exhibiting brilliant iridescence, hollow melanosomes are absent in many families containing brilliant iridescent species, such as Nectariniidae, Paradisidae and Columbidae. Still, the occurrence of a hollow modification is phylogenetically widespread. The 12 families with a hollow modification belong to 10 different orders (Galliformes, Coraciiformes, Passeriformes, Bucerotiformes, Trogoniformes, Cuculiformes, Pelecaniformes, Caprimulgiformes, Piciformes and Ciconiiformes), which suggests that the genetic changes associated with producing a hollow melanosome are either likely to occur or highly conserved in birds. A more comprehensive phylogenetic analysis will be required to determine how many times hollow melanosomes have evolved independently in birds, but our study gives a likely minimal estimate of 10 independent origins of this modification.

#### Platelet shape

We classified structures as “platelet-shaped” if they diverged from a circular cross-section. The degree of divergence varies, resulting in platelets with a range of eccentricities. Unfortunately, with few exceptions, the studies surveyed did not include measurements of the width of platelets, preventing us from quantifying and exploring the 3D shape of platelets (e.g., eccentricity). We did not find support for the hypothesis that platelets allow birds to incorporate a greater number of layers in the iridescent structure—there was no significant difference between number of layers in structures with platelets compared to rods (phylogenetic ANOVA, F(1, 220)= 21.88, p= 0.321).

Platelets are present in 11 bird families, or 31% of all families represented in the feather iridescence database (Figure 3B). This is a very similar to the frequency of the hollow modification. In fact, many of the families that have evolved a hollow modification have also evolved platelets. In some cases the modifications have evolved in combination, producing hollow platelets—but in other cases solid platelets and hollow rods have evolved separately within a family. Only Nectariniidae, Hirundinidae, Hemiprocnidae, Apodidae and Psophiidae have evolved platelet shapes but never hollow forms (with the caveat that this may change with increased sampling). As with hollowness, platelets are present in many but not all families with brilliant iridescence. For example, platelets are absent in Paradisidae, Phasanidae and Columbidae. Nevertheless, platelets are widely distributed across birds; they are present in 7 different orders (Passeriformes, Pelecaniformes, Caprimulgiformes, Trogoniformes, Gruiformes, Piciformes, Cuculiformes).

#### Evolution of multiple modifications

We hypothesized that hollow and platelet modifications are in fact different mechanisms for achieving thin melanin layers. This is supported by the fact that hollowness and platelet-shaped melanosomes always have thin melanin layers – there are no platelets or hollow forms with melanin layers ≤190nm. However, five bird families have evolved all three modifications: thin melanin layers, hollowness and platelet shape (Trochilidae, Trogonidae, Sturnidae, Galbulidae and Threskiornithidae, Figure 3B). If hollowness and platelet shape are alternative ways to achieve thin melanin layers, then why have some birds evolved both? The repeated evolution of hollow platelets suggests that at least one modification carries some additional functional value. For example, hollowness may in itself also increase the brightness of colors. Though it is possible that both modifications evolved together due to a shared mechanistic path, rather than due to some adaptive benefit, this is unlikely because species in each order with hollow platelets have close relatives with solid platelets, solid rods and/or hollow rods (Figure 3A). Thus, there does not appear to be a strong constraint on evolving particular modifications together, since each modification exist in isolation.

### 3) Optical consequences of modified melanosomes

To understand how each melanosome modification affects color production, we simulated light reflection from different structures using optical modelling. We generated 4500 unique structures which varied systematically in structural parameters (including diameter of melanosomes, lattice spacing, hollowness and platelet shape; see full model description in Methods). The parameter ranges used to generate the structures were derived from the known ranges reported in the feather iridescence database. Thus, although the simulated structures are hypothetical, they represent a realistic approximation of the structural variation that could exist, while allowing us to standardize parameters that could bias comparisons in real structures. For example, we modeled all simulated structures as photonic crystals with four layers, while real structures include single-layered structures as well as photonic crystals with varying numbers of layers, which would affect the brightness and saturation of colors independent of melanosome types.

We modeled the simulated reflectance spectra in avian color space to estimate color saturation and diversity in a manner that is relevant to bird color perception. The avian tetrahedral color space represents all the colors a bird can theoretically perceive (Endler and Mielke, 2005; Goldsmith, 1990; Stoddard and Prum, 2008). Reflectance spectra can be represented in tetrahedral avian color space as a function of how they would stimulate a bird’s four color cone types. Once reflectance spectra are mapped in avian color space, we can extract values of saturation (distance to the achromatic center of the tetrahedron) and color diversity (mean Euclidean distance between all points, and number of voxels occupied, see Methods for details). To quantify the brightness of a spectrum, we used two measures: (1) peak reflection (% reflectance at the wavelength of maximum reflectance); and 2) estimated stimulation of the avian double cones, which may play a role in achromatic perception (Hart, 2001; Jones and Osorio, 2004). We refer to both metrics as “brightness” for convenience; the term luminance is often used to describe the perception of signal intensity (here modeled using the avian double cones). Together, these metrics give a good representation of the saturation, color diversity and brightness of simulated reflectance spectra, where saturation and brightness together describe the brilliance.

Optical modeling revealed that thick solid rods are severely constrained in color diversity (Figure 6A). The simulated spectra are clustered towards the center of the tetrahedron, which means that they are producing colors of low saturation. In known feather nanostructures, thick solid rods are almost exclusively found in single-layered structures, which produce colors of low saturation and brightness. In theory, low color saturation and brightness could be due to the single-layered structure, as opposed to the melanosome type. However, we modeled all structures with four layers, suggesting that it is the thick solid rods themselves—and not the number of layers—that limits color production. In other words, producing saturated colors is not possible with thick solid melanosomes, irrespective of whether the structure is single-layered or a photonic crystal.

**Figure 6.**
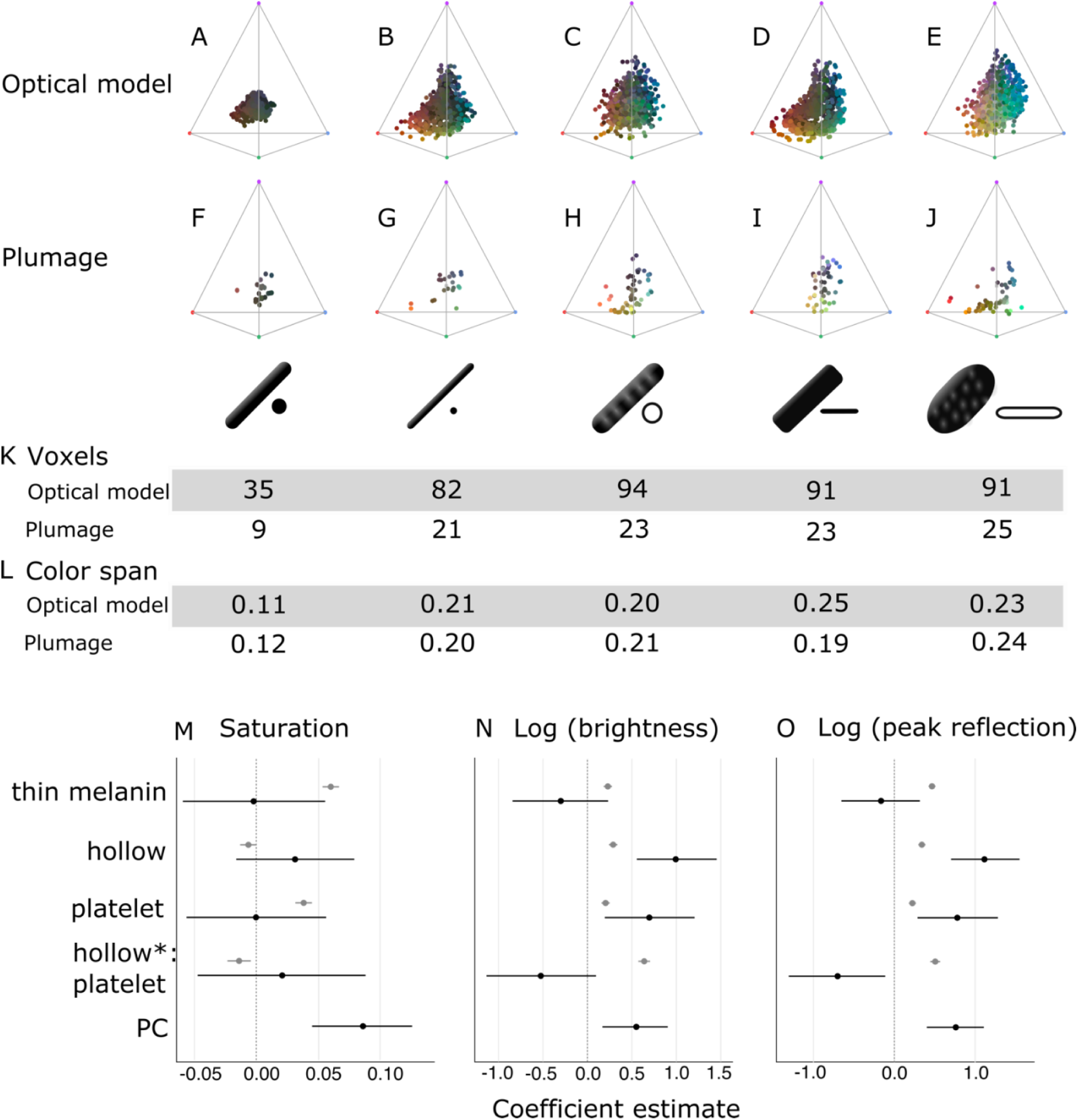
Optical effects of different melanosome modifications, as predicted by an optical model and found in empirical plumage analysis. A-J show the color diversity for structures with each type of melanosome represented in an avian tetrahedral color space (optical model, A-E; plumage data F-J). Statistics for color diversity are presented in terms of the number of occupied voxels (K) and mean color span (L) for both data sets. Thick solid rods produce colors of substantially lower diversity and saturation (A, F) than all melanosome types with thin melanin layers (B-E, G-J). In contrast, hollowness or platelet shape does not affect color diversity notably (C-E, H-J). M-O depict the estimates for the effects of each melanosome modification on saturation (M), log (brightness) (N) and log (peak reflection) (O), as predicted by linear models. The parameter PC described the variation explained by having a photonic crystal, which was used to control for variation in plumage data (see Results §4). Grey points show coefficient estimates for a model based on optical model simulations, and black dots the posterior coefficient estimates for a model based on the plumage data. Horizontal lines show 95% confidence intervals for estimates.

In contrast, all four derived melanosome types with thin melanin layers produce a large range of saturated colors (Figure 6B-E). Color diversity (voxel occupation and color span) is very similar for the four derived types, suggesting that melanin thickness is the most important modification to achieve saturated and varied colors (Figure 6K-L), which is the only modification they all share.

To explore the effect of thin melanin layers, hollowness and platelet shape on color properties in detail, we constructed linear models with melanosome modifications as binary predictors and saturation, brightness, and peak reflectance as responses. This allowed us to separate effects for each modification, which are combined in many melanosome types (for example, hollow rods are both hollow and have thin melanin layers). In agreement with the results for color space occupation, melanin layer thickness explained the greatest amount of variation in saturation in our linear model (Figure 6M). A positive effect was also seen for a platelet shape, which suggests that solid platelets produce colors of the highest saturation. Small losses in saturation are incurred from incorporating hollowness, as can be seen from the negative coefficients of the variable hollowness and the interaction term hollowness×platelet shape (describing hollow platelets).

The linear model showed that all modifications increase brightness. This effect is strongest for the interaction of hollowness and platelet shape (Figure 6N). Thus, the optical model predicts that hollow platelets produce the brightest colors. This effect likely arises from a lowered refractive index of melanosome layers with hollow platelets, which have a lower melanin-to-air ratio than layers built with hollow rods. However, this effect may be considerably weaker in real structures, where hollow platelets often have an internal honeycomb-like structure of melanin (Figure 2E), which would make the effective refractive index closer to that of hollow rods. Thin rods, hollow rods and solid platelets produce colors that are less bright than those of hollow platelets but similarly bright to one another. The linear model for peak reflection yielded similar results to those obtained for brightness (Figure 6O). These results indicate that evolving a thin melanin layer thickness is the single most important factor for dramatically increasing color diversity and saturation while simultaneously increasing brightness. When this effect is accounted for, a platelet-shape has a similar but weaker effect on saturation and brightness and hollowness only increases brightness further.

### 4) Testing predictions in real plumage data

Next, we investigated whether we could recover the same patterns in the iridescent plumage of birds with different nanostructures. We collected spectral data from 80 species that were represented in the feather iridescence database and possessed known melanosome types.

In agreement with the optical model results, the color diversity of structures with thick solid rods was low, almost half of that found in structures with thin melanin layers (Figure 6F). Moreover—mirroring the results in our optical model simulations—thin solid rods, hollow rods, solid platelets and hollow platelets are all nearly equal in color diversity (Figure 6K-L). However, some differences are noteworthy. In contrast to other melanosome types, solid platelets do not produce any saturated red colors. This is unlikely to be due to any inherent developmental or physical constraint, since our optical model simulations—based on realistic melanosome properties, including size—indicate that solid platelets can clearly occupy this area of color space (Figure 6D). Rather, this effect may be a consequence of phylogenetic bias, as the majority of species with solid platelets in our data set are sunbirds (family Nectariniidae), a group that uses carotenoid pigments—rather than structural colors— for red plumage coloration.

To further explore how thin melanin layers, hollowness and a platelet shape affect saturation, brightness and peak reflectance, we fitted generalized linear mixed models using Bayesian methods that allowed us to account for multiple measurements within a species (i.e., we obtained two reflectance measurements per plumage patch per species). In contrast to our optical model simulations, melanosomes in the real plumage patches we measured were arrayed in a variable number of layers. Since having many layers is known to increase brightness and saturation of colors, we added a parameter to control for this effect. The binary parameter “PC” (photonic crystal) described whether a structure contained a single layer of melanosomes (not a photonic crystal), or several repeating layers of melanosomes (photonic crystal).

In this linear model, there were no significant effects of either platelet shape or hollowness on saturation (Figure 6M). Thus, we did not find support for the optical model prediction that solid platelets produce more saturated colors. We also did not find a significant positive effect of thin melanin layers (Figure 6M), in contrast to our findings with the optical model. However, this is likely due to the fact that—in real plumage—thick solid rods are exclusively found in single-layered structures. Thus, the model can only compare color for *single-layered structures* with thick and thin melanin layers. The entire difference in saturation between structures with thin and thick melanin layers (if it exists) will therefore be captured by the PC parameter. Thus, these results reveal that melanin thickness in single-layered structures does not affect saturation. While the plumage data cannot directly tell us the effect of thin melanin layers in a photonic crystal, we do show that single-layered structures with thick solid rods (plumage data) are just as constrained in color diversity as the simulated colors generated by photonic crystals with thick solid rods (optical model) (Figure 6A, F). Thus, the optical model predicts that adding more layers with thick solid rods would not increase color saturation or diversity. This is very likely the reason that no such structures exist. In single-layered structures on the other hand, the thickness of melanin layers is typically irrelevant because interference occurs between light reflected from the top and bottom of the overlying keratin cortex (Doucet, 2006).

In terms of brightness and peak reflectance, the plumage data compare to the optical model simulations in interesting ways. In agreement with the optical model, the linear model revealed a significant positive effect of hollowness and platelet shape on the brightness and peak reflectance of colors (Figure 6N-O). However, we did not see a large positive effect of hollow platelets in the empirical data. In fact, this parameter has a negative effect, which is significant for peak reflectance (Figure 6O). This discrepancy may be due to the fact that–in the real plumage structures measured—hollow platelets tended to be arranged in relatively few layers. Our sample of structures with solid platelets consisted almost entirely of different species of sunbirds (Family Nectariniidae), which exhibit 5-8 layers (Durrer, 1962), while the sample for hollow platelets contained several groups with fewer layers (e.g., *Priotelus* and *Apaloderma*, (Durrer, 1977)). We could not control for this because the number of layers is not known in many of the structures we sampled; instead, we only included a parameter to indicate if a structure was a photonic crystal or not. We can, however, compare the brightness of *single-layered* structures with hollow platelets versus solid platelets. This comparison shows that the hollow platelets produce brighter colors (phylogenetic ANOVA, F(1,34)=12.10, p=0.034, df=1). Thus, the general conclusion that hollowness increases brightness is well supported, although this advantage is likely to diminish with increasing number of layers in the structure. Reflection from a multilayer with melanin and keratin becomes saturated at >9 layers (Land, 1972), thus it is likely that the greatest advantage of hollowness is gained for structures with nine layers or less.

In summary, the plumage data support the general conclusions that thin melanin layers are critically important for producing diverse and brilliant colors, while hollowness and platelet shape are less crucial. We observe a near doubling of color diversity for real plumage structures with thin melanin layers compared to structures with thick solid rods, consistent with results of the optical model. While the plumage data alone cannot prove that this difference is driven by thin melanin layers rather than simply the PC parameter by itself, our optical models exclude this possibility (see Figure 6A for a simulation of photonic crystals with thick solid rods). Hollowness and a platelet shape increase the brightness of colors further, in agreement with the optical model.

## Discussion

Brilliant iridescence has been linked to the evolution of different melanosome modifications, most notably hollowness and a platelet shape (Eliason et al., 2013; Maia et al., 2013b), but how these modifications affect color production has not been evaluated in a unified framework. Here, we have taken a broad approach comparing all five melanosome types found in iridescent feathers to uncover general design principles governing the production of brilliant iridescence. We find that the most important modification to increase brilliance is not hollowness or a platelet shape *per se*, but rather a third modification that unites all melanosomes found in brilliant iridescent structures: thin melanin layers. Specifically, we show that melanosomes in brilliant structures have converged on a melanin layer thickness of approximately 40-200nm (Figure 5), which is the theoretical optimal thickness to produce first-order interference peaks in the bird-visible spectrum. Our optical simulations and empirical data demonstrate that this modification alone nearly doubles color diversity (Figure 6A-L) and simultaneously increases saturation and brightness (Figure 6M-O). In contrast, hollowness and platelet shape on their own only contribute to increased brightness.

Our results have interesting implications for the evolution of brilliant iridescent structures in birds. We show that two key optical innovations are required: a photonic crystal (multiple periodic layers of melanosomes) and 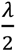 thick melanin layers. Indeed, Durrer, 1977 observed that these two features were common to the brilliant structures he studied, and here we validate the importance of his observation with color measurements and optical modelling. Since photonic crystals with all four melanosome types found in brilliant iridescent structures have similar optical qualities, this suggest that variability in melanosome type may be strongly influenced by historical factors, as opposed to particular types being associated with specific optical functions. Thus, the reason that sunbirds (Nectariniidae) produce brilliant iridescence with solid platelets while hummingbirds (Trochilidae) mainly use hollow platelets (Figure 3A) is likely related to variation in evolutionary history rather than to variation in selection for different optical properties. Supporting this interpretation is the fact that diverse photonic crystals in birds often have independent evolutionary origins. In Galliformes, some families have photonic crystals with thin rods, and others have photonic crystals with hollow rods (Figure 3A), but these different structures have almost certainly evolved from an ancestor with a non-iridescent or single-layered structure rather than a photonic crystal (Gammie, 2013). Similarly in Sturnidae, photonic crystals with hollow rods in *Cinnyricinclus* and photonic crystals with hollow platelets in *Lamprotornis* likely evolved independently from non-iridescent structures (Figure 3A, Durrer, 1970; Maia et al., 2013b).

Yet, in some groups, melanosome type is highly variable within the same genus, or even within the same species (interpatch variability). In the birds-of-paradise (Paradisaeidae), who typically display photonic crystals with thin rods, two species (*Paradisaea rubra* and *Parotia lawesii*) are known to have evolved large rods with a porous interior (Durrer, 1977; Stavenga et al., 2015). In *Parotia lawesii*, other iridescent patches contain structures with thin solid rods, proving interpatch variability in melanosome type. Hummingbirds, whose iridescent structures are typically built with hollow platelets, can also exhibit interpatch variability in melanosome type. Some patches may contain a structure with solid platelets, or even mixed structures with both hollow and solid platelets (Gruson et al., 2019). It is notable that the only known examples of interpatch variability in melanosome type comes from the birds-of-paradise and hummingbirds—groups that are known to have exceptionally high rates of color evolution (Eliason et al., 2013; Ligon et al., 2018; Parra, 2010). One hypothesis to explain this variation could be that modifications in hollowness/platelet-shape tune the brightness of some patches (Figure 6N-O). However, this seems unlikely. Both birds-of-paradise and hummingbirds typically have >9 melanosome layers in their iridescent structures, which already achieves nearly 100% reflectance irrespective of melanosome type. Moreover, our results suggest that there would be little or no difference in brightness between structures with solid platelets and hollow platelets (only between thin rods and hollow and/or platelet-shaped melanosomes), which is the variability we see in hummingbirds. Indeed, Gruson et al., 2019 found color production to be similar among patches with different melanosome types in hummingbirds. We speculate that high variability in melanosome type in hummingbirds and birds-of-paradise is not related to general optical benefits of specific melanosome types or modifications, but rather to general high rates of color change in these groups (Eliason et al., 2020; Parra, 2010). Our optical modeling results (Figure 6B-E) show that there are multiple ways to reach the same areas of color space—using different melanosome types. It is possible that a change in melanosome type may be the fastest route to a new area of color space, even though the same color shift could in theory be produced with adjusting the size of the original melanosome type. This idea is hard to test with our current very limited understanding of the genetics of iridescent structures, but it does predict that groups with high variation in melanosome type have a greater standing variation in genetic traits associated with different melanosome types. It also predicts that patches with higher rates of color evolution should also have greater variability in melanosome type.

However, we cannot fully exclude hypotheses based on general adaptive explanations tied to melanosome type to explain interpatch variability in melanosome type. We did not investigate differences in angular variation of color for different types of structures, or potential non-signaling functions such as differences in mechanical properties of the barbule (Burtt Jr., 1979) and microbial resistance (Goldstein et al., 2004). These topics are promising avenues of future research. Nevertheless, it is clear that we need to understand how brilliant structures evolve to resolve fully the mystery of their structural diversity. To our knowledge, no general models have been proposed to explain how photonic crystals with modified melanosomes evolve from more simple, single-layered structures (but see discussion by Durrer, 1977, 1970). We can use the insights derived from our study to propose two hypothetical routes to brilliant iridescence.

Brilliant iridescent structures likely originate from single-layered structures with thick solid rods (Maia et al., 2012; Shawkey et al., 2006). To achieve brilliant iridescent colors, such a structure must evolve to incorporate a photonic crystal-like organization of melanosomes—and the melanosomes must have thin melanin layers. However, our results showed that either of these changes on their own does not increase color saturation or brightness. This leads to an interesting problem, where only the two adaptations *together* produce a great advantage in brilliance. How could such a structure evolve? We suggest two plausible evolutionary routes by which this conundrum can be solved (Figure 7).

**Figure 7.**
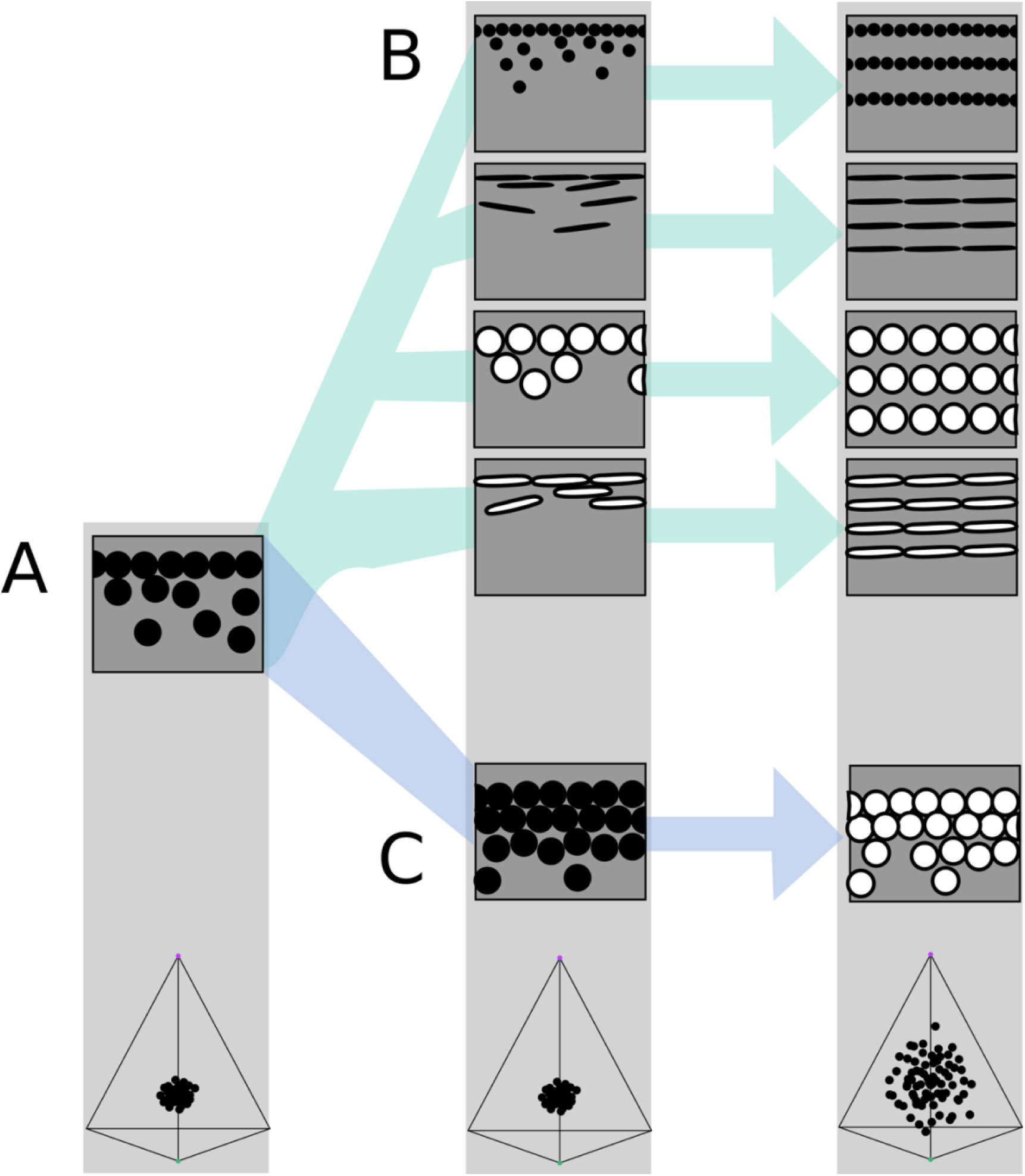
Hypothetical evolutionary paths to brilliant iridescence. Grey squares depict schematics of barbule cross-sections, showing the iridescent nanostructures within, while the tetrahedra below show hypothetical color diversity for each evolutionary “step”, represented in avian color space. A) Assumed ancestral state for iridescent structures - a single-layered structure with thick solid rods. Note that a layer refers to a *continuous* layer of melanosomes; scattered or disorganized melanosomes often seen below a continuous single layer do not constitute additional layers. From this state, structures may either immediately evolve modified melanosomes in a single-layered structure (B) or first evolve multilayered, hexagonal structuring of thick solid rods (C). Both of these states are expected to give a negligible advantage in terms of color saturation and diversity, as seen in the hypothetical color spaces corresponding to each stage (bottom). We argue that path B might initially be driven by selection for brighter colors, while path C could form spontaneously from higher concentrations of melanosomes in the barbule. Both paths can then evolve towards more brilliant forms (multilayers in B, modified melanosomes with thin melanin layers in C) which will drastically expand the possible color diversity.

In the first route, modified melanosomes with thin melanin layers evolve for reasons unrelated to color saturation (Figure 7B), perhaps to enhance brightness. Hollow and platelet-shaped modifications may evolve initially to produce brighter colors, while thin solid rods have been hypothesized to facilitate formation of thin-film structures through their elongate shape (Maia et al., 2012). Once evolved, melanosomes with thin melanin layers allow for the evolution of photonic crystals, since such structures would produce brighter and more saturated colors. The second route to brilliant iridescence involves the spontaneous formation of a photonic crystal from a single-layered structure, which then selects for modified melanosomes with thin melanin layers (Figure 7C). In many single-layered structures, a discontinuous second layer can be seen beneath the top layer, where melanosomes are packed hexagonally (e.g., Figure 2A). This likely provides a more mechanically stable configuration during barbule development (as suggested by Eliason et al., 2013). It is easy to see how the evolution of hollowness in such a structure would lead to the production of brilliant iridescence.

The feather iridescence database gives some support to both of these hypothetical routes. Single-layered structures with modified melanosomes are relatively common (Figure 5), suggesting that this may be a likely first step towards more complex structures. Similarly, hexagonally arranged photonic crystals with hollow rods are common in many groups (Galliformes, Trogoniformes), which also contain taxa with single-layered structures with thick solid rods. However, very few clades are sampled in sufficient detail to draw inferences about the transitions between different structures. To test our hypotheses, careful characterization of nanostructures in a group with repeated transitions to brilliant iridescence is needed. Such a study could also lay the groundwork for exploring the genetic regulation of iridescent structures, an area of research in its infancy (Saranathan and Finet, 2020).

By investigating the evolution and optical properties of brilliant iridescent feather nanostructures spanning 15 avian orders, we have identified some features common to iridescent nanostructure design and some features that are likely to result from differences in evolutionary history. The key feature uniting melanosomes in brilliant iridescent structures is the presence of thin (40-200nm) melanin layers, which tunes a photonic crystal optimally to produce bright and saturated colors in the bird-visible spectrum. We suggest that much of the diversity in melanosome type in brilliant iridescent structures - such as the prevalence of solid platelets in sunbirds but hollow platelets in hummingbirds - could be explained by differences in evolutionary history, since different melanosome types offer alternative routes to producing thin melanin layers. However, the large scale-patterns uncovered in this study are only a first step towards gaining a deeper understanding of how these dazzling structures have evolved. We propose two likely evolutionary routes, which could be tested further by careful study of a clade with repeated transitions to brilliant iridescence. A focus of future studies should be to explore the evolutionary steps associated with the evolution of brilliant iridescence, and ultimately to tie these steps to genetic changes.

## Materials and methods

### Building the feather iridescence database

We surveyed the literature for microscopy studies of iridescent feathers using two complementary approaches. For studies published earlier than 2006, we used the references in Prum, 2006 and Durrer, 1977 as a starting point. For later publications, we used Google Scholar to search for articles containing the terms “iridescence” and “feather”. We then extracted the following information from each study (where available, or possible to infer from redundant measurements): melanosome arrangement (single-layered, photonic crystal), melanosome type (i.e., solid rod, hollow rod, solid platelet or hollow platelet), melanosome diameter (d_melsom_), lattice spacing (a), the number of melanosome layers (n), diameter of hollow interior (if present, d_air_), thickness of keratin layers (ks), thickness of melanin layers (mt; for solid forms mt= d_melsom_, for hollow forms mt= (d_melsom_ - d_air_)/2), cortex thickness (c), the patch from which the studied feather originated, and the color of the feather. A schematic of all measurements is shown in Figure 8. With few exceptions, most studies sampled only a single iridescent patch from each species. This is based on the assumption that iridescent nanostructures are similar in all iridescent patches in a species, which seems to be true in most species but not all; hummingbirds and birds-of-paradise are the only known exceptions (Durrer, 1977; Gruson et al., 2019).

**Figure 8.**
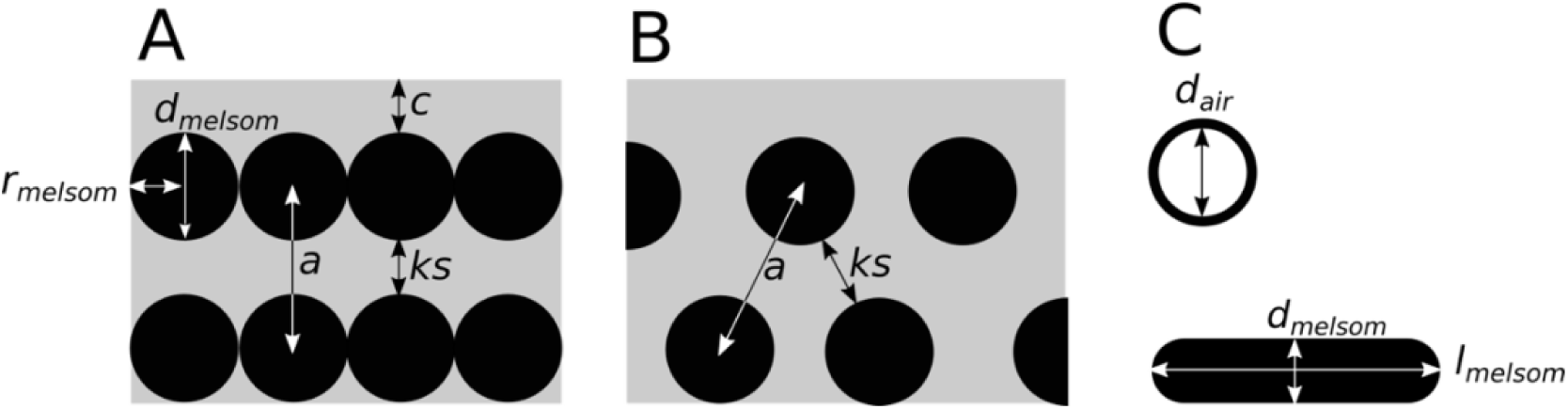
Definitions of parameters used in the study, shown in schematics of cross-sections of iridescent structures: (A) laminar photonic crystal (multilayer); (B) hexagonal photonic crystal; and (C) isolated hollow and flat melanosomes. In (A), d_melsom_: diameter of melanosome (shortest axis in flat melanosomes), r_mel_: radius of melanosome (d_melsom_ /2), c: thickness of keratin cortex, a: lattice spacing (center-center distance between melanosomes), ks: keratin spacing (thickness of keratin layer between melanosomes at the thinnest point). In (B) keratin spacing (ks) and lattice spacing (a) is shown for a hexagonal photonic crystal. In (C), d_air_: diameter of internal air pockets (shortest axis of air pockets in hollow platelets), l_melsom_: width of platelets. Melanin layer thickness is defined as d_melsom_ for solid forms, and (d_melsom_ - d_air_)/2) for hollow forms.

For a small number of records (n=17), we produced new measurements of iridescent structures using transmission electron microscope images previously collected by Nordén et al., 2019. Images were measured using the program ImageJ (Abràmoff et al., 2004). All images used for new measurements are included in the Supplementary Information.

In total, our database covers 46 studies from 1952-2019 and 306 unique species, across 35 families and 15 orders (37% of total orders and 20% of total families in Aves).

### Phylogeny

We used the phylogenies of Jetz et al., 2012, which are based on a Hackett et al., 2008 backbone, to construct a tree including all the species in the feather iridescence database and the species from the Li et al., 2012 dataset. We sampled 1000 pruned trees from the tree distribution available at birdtree.org and then constructed a 50% consensus tree from this distribution. Branch lengths were calculated using the “consensus.edge” function in the R package *phytools* (Revell, 2012). This tree was then pruned as necessary for different analyses.

### Optical modeling

We modeled the reflectance from iridescent feather structures using the software package MIT Electromagnetic Equation Propagation (MEEP) (Oskooi et al., 2010). Simulations were performed in one unit cell, with an absorbing perfectly matched layer in the x-direction, and periodic boundaries in the y-direction. Resolution was set to 80 pixels/um, which gives 12 sampling points for one 300nm wave in the material with the highest refraction index (melanin). We set the extinction coefficient (k) of melanin to 0.1, the refractive index (n) of keratin to 1.56, and the refractive index of melanin to 2. In reality n and k for most materials vary over the spectrum, and these values are an approximation based on published values (Brink and van der Berg, 2004; Stavenga et al., 2015). However, we do not expect small differences in these parameters to alter the larger patterns we have described (though they may alter the exact hue and reflection of a particular structure). The extinction coefficient for keratin is likely to be low (k=0.03, Brink and van der Berg, 2004) and was omitted (set to 0).

The structural parameters varied in the model were melanosome diameter (d_melsom_), relative hollowness (d_air_/d_melsom_), flatness (l_melsom_/d_melsom_), relative lattice spacing (r_melsom_/a), and cortex thickness (c, Figure 8). We set the ranges for parameters related to melanosome shape to match the known ranges for each melanosome type, extracted from the feather iridescence database. Lattice spacing and cortex thickness were varied in the same way for all melanosome types (the overall range in the feather iridescence database), and number of layers was fixed to 4 (the median in the feather iridescence database). For structures with rods, we modeled structures with a hexagonal packing in addition to the standard laminar configuration (Figure 8B and A respectively) to represent the diversity present in real structures. Although a square configuration also exists, we did not model this since it has only been recorded in a single genus, the peafowls (*Pavo*). Table 1 gives a detailed overview of the model settings for each melanosome type. Notice that the melanosome diameter of solid forms is varied in 30 steps, while the diameter of hollow forms is only varied in 10 steps. The thickness of melanin layers is important for determining hue, and hollow forms have two parameters that adjust this value (diameter and hollowness), while solid forms have only one (diameter). To avoid a bias towards greater hue variability in hollow forms due to this effect, we allowed the diameter of solid forms to vary in an equal number of steps as the combined effect of diameter and hollowness in hollow forms (10×3=30).

**Table 1.**
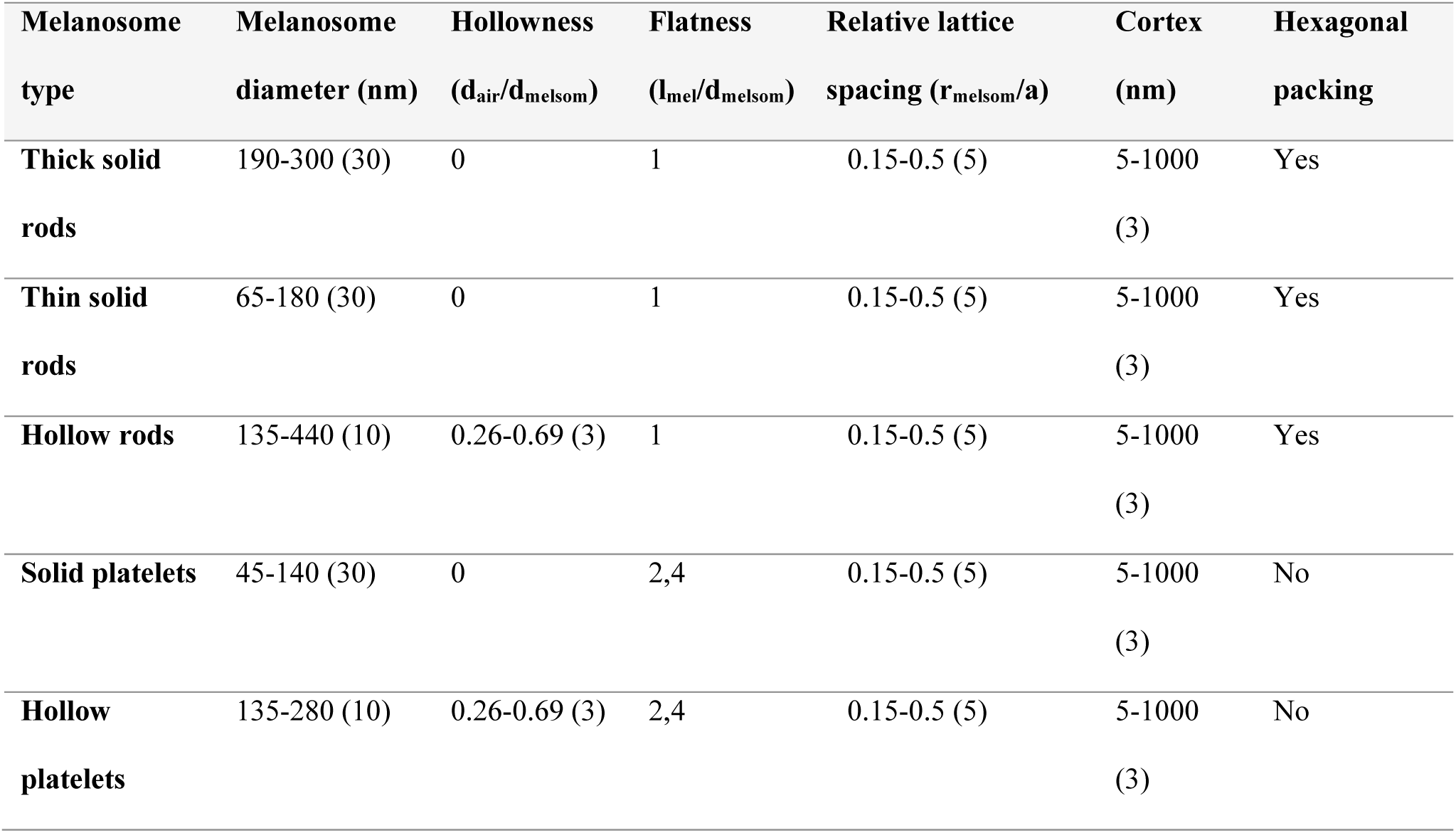
Model parameter ranges for each melanosome type. The values reported in parentheses are the number of evenly spaced steps with which the parameter was varied. For each melanosome type, we simulated 900 unique structural configurations.

| Melanosome type | Melanosome diameter (nm) | Hollowness ($d_{\text{air}}/d_{\text{melosom}}$) | Flatness ($l_{\text{mel}}/d_{\text{melosom}}$) | Relative lattice spacing ($r_{\text{melosom}}/a$) | Cortex (nm) | Hexagonal packing |
| --- | --- | --- | --- | --- | --- | --- |
| Thick solid rods | 190-300 (30) | 0 | 1 | 0.15-0.5 (5) | 5-1000 (3) | Yes |
| Thin solid rods | 65-180 (30) | 0 | 1 | 0.15-0.5 (5) | 5-1000 (3) | Yes |
| Hollow rods | 135-440 (10) | 0.26-0.69 (3) | 1 | 0.15-0.5 (5) | 5-1000 (3) | Yes |
| Solid platelets | 45-140 (30) | 0 | 2,4 | 0.15-0.5 (5) | 5-1000 (3) | No |
| Hollow platelets | 135-280 (10) | 0.26-0.69 (3) | 2,4 | 0.15-0.5 (5) | 5-1000 (3) | No |

In total, we ran 4500 simulations, with 900 simulations for each melanosome type.

### Plumage measurements and spectral analysis

We collected spectral measurements of 80 bird species (across 13 orders) for which nanostructures were already known (see references in the feather iridescence database), housed in the American Natural History Museum, New York. Spectral measurements were taken directly on the specimen following standard procedures (Andersson and Prager, 2006). Briefly, we used a USB4000 spectrophotometer and a PX-2 xenon light source (Ocean Optics, Dunedin, FL, USA). We measured color over a range of angles (15°-135 °) using a goniometer, keeping the light source fixed at 75° (Figure S1). Two individuals were used for each species, and all iridescent patches with different color (as perceived by human vision) were measured. In total, 120 unique patches were measured.

Spectra were analyzed in R v. 3.6.1 (R Core team, 2019) using the package *pavo* (Maia et al., 2013a). All spectral data were first smoothed to remove noise, using locally weighted smoothing (LOESS) and a smoothing parameter of 0.2. We then extracted the spectra with maximum total brightness (area under the curve) for each patch. The variability between individuals of each species was assessed using pairwise distances in tetrahedral color space. If the patch measurements for the two individuals were very different in terms of color (separated by >0.1 Euclidean distance in color space), we inspected the spectral measurements to identify possible inaccurate readings. Eight spectra were removed from the data set after this process, leaving a total of 232 spectra used for analysis.

### Calculation of color variables used in analysis

To compare color diversity and color properties of different structures, we focused on five variables: 1) the number of voxels occupied in avian color space, 2) mean color distance in avian color space, 3) color saturation, 4) stimulation of the avian doble-cone (brightness) and 5) peak reflectance. These variables describe color diversity (1-2), color purity (3), perceptual brightness (4), and objective brightness (5) respectively. Peak reflectance is simply the maximum reflectance from each spectrum. Perceptual brightness was modeled as the photon catch from a chicken double cone (*Gallus gallus*, built-in data in the *pavo* package; see details above), since current evidence suggests that the double cones mediate achromatic/brightness perception in birds (Hart, 2001; Jones and Osorio, 2004). Saturation and color diversity were based on modelling spectra in avian color space (Stoddard and Prum, 2008). This space represents all the colors a bird can theoretically perceive. Relative cone stimulation was calculated from photon cone catches using cone sensitivity functions in *pavo*. Bird species vary in their ultraviolet spectral sensitivity; some species have a VS (violet-sensitive) cone type that is maximally sensitive in the violet range while others have a UVS (ultraviolet-sensitive) cone type that is maximally sensitive in the ultraviolet range. Because we modeled plumage colors across many phylogenetic groups, we used the sensitivity curves in pavo for an “average UVS” (λ_max_ = 372nm) and “average VS” (λ_max_ = 416nm) type system. Since results in general were similar for a UVS- and VS-type system, we only include analyses based on a VS-type visual model (summary statistics for a UVS-type cone can be found in Table S5-6), which is the ancestral condition in birds (Ödeen and Håstad, 2003).

Saturation in tetrahedral color space is simply the distance from the center of the tetrahedron (r vector, as defined by Stoddard and Prum, 2008). For number of voxels occupied, we followed the approach of Delhey, 2015. The tetrahedral color space is divided into 3D pixels (voxels), and then the number of voxels that have at least one data point are counted. The resolution of raster cells was set to 0.1, which gives a total of 236 voxels in tetrahedral color space. Mean color span is a measure of the spread of samples in color space and is calculated as the mean of pairwise Euclidean distances between all samples. This measure is more robust to sample size differences than voxel occupation, which makes it better suited to compare our plumage data.

### Statistical analysis

To compare iridescent structures recorded in the feather iridescence database (thickness of melanin layers, diameter of interior hollowness, number of layers), we applied simulations-based phylogenetic analyses of variance (ANOVA), as described by Garland et al., 1993 using the R package *phytools* (Revell, 2012). Since this function assumes Brownian evolution of traits, we measured phylogenetic signal in the traits tested to confirm that this assumption was not violated. All traits tested recorded a high and significant lambda (Table S2). To clarify relationships between groups, we also performed phylogenetic pairwise t-tests where necessary (using the R package *phytools* (Revell, 2012), Table S1). Species which had more than one entry in the database (for example, from multiple studies or multiple patches) were averaged before analysis. For comparison, we also include melanosome diameters from black feathers in some analyses. These data were taken from Li et al., 2012.

We performed a test for multimodality to assess whether solid rods show a binary distribution, following the method described by Fisher and Marron, 2001, which is implemented in the R package *modetest* (Ameijeiras-Alonso et al., 2018).

To explore how melanosome modifications affect color production, we fitted separate linear models with response variables saturation, brightness, and peak reflectance. Brightness and peak reflectance were log-transformed before inclusion into the models to achieve normally distributed residuals. We used binary predictors to describe absence/presence of the three melanosome modifications: thin melanin layers (≤190nm), hollowness and platelet shape. We also added the interaction term {hollowness×platelet}, since the optical effect of hollow platelets is not expected to be simply the addition of hollowness and platelet shape. This is because hollow platelets lower the refractive index of melanosome layers by having relatively less melanin in each layer. This property only applies to melanosomes which have both modifications simultaneously. Note that since we have included an interaction term, the variables hollow and platelet are only describing a situation where the interaction is zero, *i.e*. for hollow rods and solid platelets respectively.

Spectral data derived from optical simulations were analyzed using multiple linear regressions with the variables described above (summary of results can be found in Table S7-S9). For plumage data, we also needed to account for phylogenetic relatedness, as well as individual variability in patch color (for each species we had measurements from two individuals). We did this using Bayesian linear mixed models, adding phylogenetic structure and patch as random factors in the model. The phylogeny used was the same as for earlier analyses but pruned to contain only the 80 species in our plumage measurements. We also added a fourth predictor: presence/absence of a photonic crystal (PC). This variable accounts for expected variation in color brightness and saturation that is explained by whether the structure has a single layer of melanosomes or several (in the optical model simulations, all structures had four layers). We used the R package *MCMCglmm* (Hadfield, 2010) to run our Bayesian model with Markov Chain Monte Carlo methods. We ran chains for each model for 50 million generations, with a sampling frequency of 500. The first 50000 generations were discarded as burnin. We used the default priors for the fixed effects and set an inverse gamma distribution prior for the variance of residuals and random factors. We checked that the analysis had reached a stable phase by visually examining trace plots and checked that autocorrelation values between parameters was low (all <0.1). We also formally tested convergence of the chain using Heidelberg’s and Welch’s convergence diagnostics (all variables passed both tests). Summary of results for each model can be found in Table S10-S12.

## Acknowledgements

We would like to thank the American Museum of Natural History (NYC) for allowing us to use the bird skin collections for plumage spectral color measurements. In particular, we thank Paul R. Sweet, collections manager for the ornithological collections, for his help. We thank Kaspar Delhey for generously sharing R code to reproduce his measure of color diversity (voxel occupation). Funding in support of this work was provided by Princeton University (M.C.S.), a Packard Fellowship for Science and Engineering (M.C.S.), the Grainger Bioinformatics Center (C.M.E.), and the National Science Foundation (Award 2029538 to M.C.S.). We would also like to thank Raphael Steiner, Jarome Ali and Merlijn Staps for helpful discussion and providing feedback on an earlier version of this manuscript.

